# Antibiotics affect migratory restlessness orientation

**DOI:** 10.1101/2020.02.14.949651

**Authors:** Yuval Werber, Eviatar Natan, Yizhar Lavner, Yoni Vortman

**Affiliations:** Hula Research Center, Department of Biotechnology, Tel-Hai College; The Aleph Lab, Oxford, England; Department of Computer Sciences, Tel Hai College; Hula Research Center, Department of Animal Sciences, Tel-Hai College

## Abstract

Magnetoreception is a sense that allows the organism to perceive and act according to different parameters of the magnetic field. This magnetic sense plays a part in many fundamental processes in various living organisms. Much effort was expended in finding the ‘magnetic sensor’ in animals. While some experiments show a role of the ophthalmic nerve in magnetic sensing, others show that effects of light on processes in the retina are involved. According to these inconclusive and puzzling findings, the scientific community has yet to reach an agreement concerning the underlying mechanism behind animal magnetic sensing. Recently, the symbiotic magnetotaxis hypothesis has been forwarded as a mechanistic explanation for the phenomenon of animal magnetoreception. It suggests a symbiotic relationship between magnetotactic bacteria (MTB) and the navigating host. Here we show that in contrast to the control group, antibiotic treatment caused a lack of clear directionality in an Emlen funnel experiment. Accordingly, the antibiotics treatment group showed a significant increase in directional variance. This effect of antibiotics on behaviors associated with animal magnetic sensing is, to the best of our knowledge, the first experimental support of the symbiotic magnetotactic hypothesis.

## Introduction

Actively motile animals need to be able to navigate their environment. Using memory, timing and external cues, organisms move deliberately and reach their designated destination. Any non-random environmental feature is a candidate for use in orientation, from chemical cues [1,2], and visual characteristics of the environment [3,4] to movement of celestial bodies [5] and seismic signals [6]. In most cases, a combination of such signals is used [7].

Navigation using Earth’s magnetic field is ubiquitous throughout the entire tree of life. It occurs in a great variety of organisms, playing a part in many natural processes, from bacterial movement to global migrations. Although magnetic sensation in navigating organisms has long been accepted as fact, its underlying mechanism in multicellular animals is still being debated [8].

The “magnetite based magnetoreception” hypothesis suggests that biogenic magnetite crystals serve as magnetic field sensors by arousing mechanosensitive protein structures which translate mechanic excitation to sensory information [8]. Alternatively, the “radical pair” hypothesis predicts that the geomagnetic field modulates the outcome of biochemical reactions by influencing the spin state of light-induced radical pairs in macromolecules on vertebrate retinas [9]. Both theories fail to provide a complete magnetoreception mechanism which is functional in a natural setting.

In the absence of convincing proof for a ‘magnetotactic sensor’, a new hypothesis was proposed. The hypothesis suggests the existence of a symbiosis between magnetotactic bacteria and magnetotactile vertebrates [10]. Magnetotactic bacteria (MTB) are gram-negative aquatic prokaryotes which sense and act upon a magnetic field [11]. MTB mineralize ferromagnetic crystals in unique organelles called magnetosomes, which are arrayed on the longitudinal body axis of the bacterium, and respond to the ambient magnetic field much like a compass needle [12]. Here, we examine the effects of antibiotics on the orientation of a magnetic-sensing migrating passerine in order to provide first experimental support for the symbiotic magnetic-sensing hypothesis. Led by a straightforward rationale, we explore whether exposure to an antibiotic substance affects orientation–related, magnetic sensing behaviors of migrating passerines, using a well-established protocol to quantify the influence of an antibiotic substance on navigation-related behaviors.

## Materials and Methods

The experiment took place during the spring of 2018 (March-May) at the Hula Research Center in Israel’s Hula Valley (33°06’43.8″N, 35°35′8.1″), a major stopover site for avian migrants on the Eurasian-African flyway every autumn and spring.

### Study animal

Eurasian reed warblers (*Acrocephalus scirpaceus*) are small, night-migrating passerines that show wide latitudinal variation in their breeding grounds and migration date [13]. The location of the Hula Valley with respect to Eurasian reed warblers’ migration route suggests that during autumn migrating individuals will show a southward directional tendency and during spring most individuals are expected to show a northward tendency. Eurasian reed warblers were caught early in the morning at the Hula Ringing Station using mist nets. We chose Eurasian reed warblers with high fat scores (≥ 3), as indicating that they are preparing to migrate soon [14]. We only took specimens with primary wing feather length of ≥ 66 mm, to make sure that they belong to populations that breed at higher latitudes and are not intending to breed in or around the valley [13], which would mean their migration has ended. Suitable specimens were put into cloth bags and taken to the research station.

#### Experimental procedure

Birds were housed in 30 cm × 23 cm × 40 cm wooden cages inside an air-conditioned container. Each cage had a wooden perch, two water-filled bottle caps, and a retractable tray for food provision. Since onset of capture birds had no view of the sky or the outside environment. The first 24 hours after capture were set as an adjustment period. Birds that were eating and did not show signs of stress by the evening of the day of capture were considered adjusted, and were given food *(Tenebrio molitor)* and water ad libitum. At later stages of the experiment, a bird that appeared stressed was immediately released. On the second day, birds were randomly divided into a control group and a treatment group, and the first dose of treatment (or water, according to group) was orally administered using a pipette. Antibiotic substance, dosage, and method of administration were chosen according to avian veterinary advice, as would be administered for treating bacterially induced symptoms in the oculonasal region. Enrofloxacin, also known as Baytril, is a standard, FDA-approved substance for treating bacterial infections in vertebrates from all groups. It is a broad-spectrum antibiotic, which we tested against MTB during June 2017 at the molecular and environmental microbiology lab in CEA, Cadarache, France, with the aid of Dr. Christopher Lefevre. Enrofloxacin proved lethal to various MTB types including various unidentified morphotypes from the research area and cultivated *Magnetospirillum magneticum* strain AMB-1 and *Magnetovibrio blakemorei* strain MV-1. One dose comprised 2 μl of the solution (or water), and each bird received four doses: one in the morning and one before sunset, for two consecutive days. On the third day after capture, half an hour after sunset, birds were removed from their cages and placed in Emlen funnels (plastic funnels with a rim diameter of 45 cm, covered by a PVC sheet), which were placed in a mesh enclosure with no view of the sky. Funnels were filmed from above using HlKvision 2.8 IR cameras (one for each funnel), connected to an HIK vision HD DVR hard drive. Recording started exactly one hour after sunset with a shot of the identity number and a compass, to indicate north for later analysis. Filming continued for 90 minutes, of which the first ten allowed for recovery, and were not analyzed. At the end of the 90-minute experiment birds were removed from the funnels and released.

#### Statistical analysis

Each bird’s average hop azimuth was transformed to its projection on both axes of the trigonometric unit circle according to [15]. We used Rayleigh’s Z test to examine whether individuals and groups displayed significant directionality. To verify that the directional data fit a unimodal distribution, we assessed the distribution using the AIC criterion, and dedicated methods for circular data, according to [16]. Comparison of the variance in directional tendencies between groups was done using Levene’s test. Activity levels were compared using two-tailed t-tests.

## Results and Discussion

When examining orientation of individuals placed in Emlen funnels, we found significant difference in orientation between the two experimental groups. Individuals from the control group showed a highly significant south-westward orientation (n = 14, Z = 8.84, P < 0.0001, Figure 1A), while individuals from the treatment group were more dispersed and showed no significant orientation (n = 14, Z = 1.33, P > 0.2, Figure 1B). According to the significant orientation of controls and lack of significant orientation of the treatment group, using Levene’s tests we show that variance of directionality was significantly lower in the control group in both the north/south and the east/west axes: (north/south axis: F = 6.5, P = 0.01, sd: control = 0.5, treatment = 0.78, Figure 2A; east/west axis: F = 7.3, P = 0.01, sd: control = 0.38, treatment = 0.6, Figure 2B). As seen by standard deviation values, controls exhibit smaller variance in all aspects. To verify that the lack of significant orientation of individuals from the treatment group is not a result of reduced activity due to exposure to antibiotics, we compared the number of hops between groups as a means of negating any non-navigation-related effects of antibiotics. Groups did not differ in the number of hops per individual (T = 0.1, n _c_ =14, n _t_ = 14, P = 0.9).

**Figure 1:**
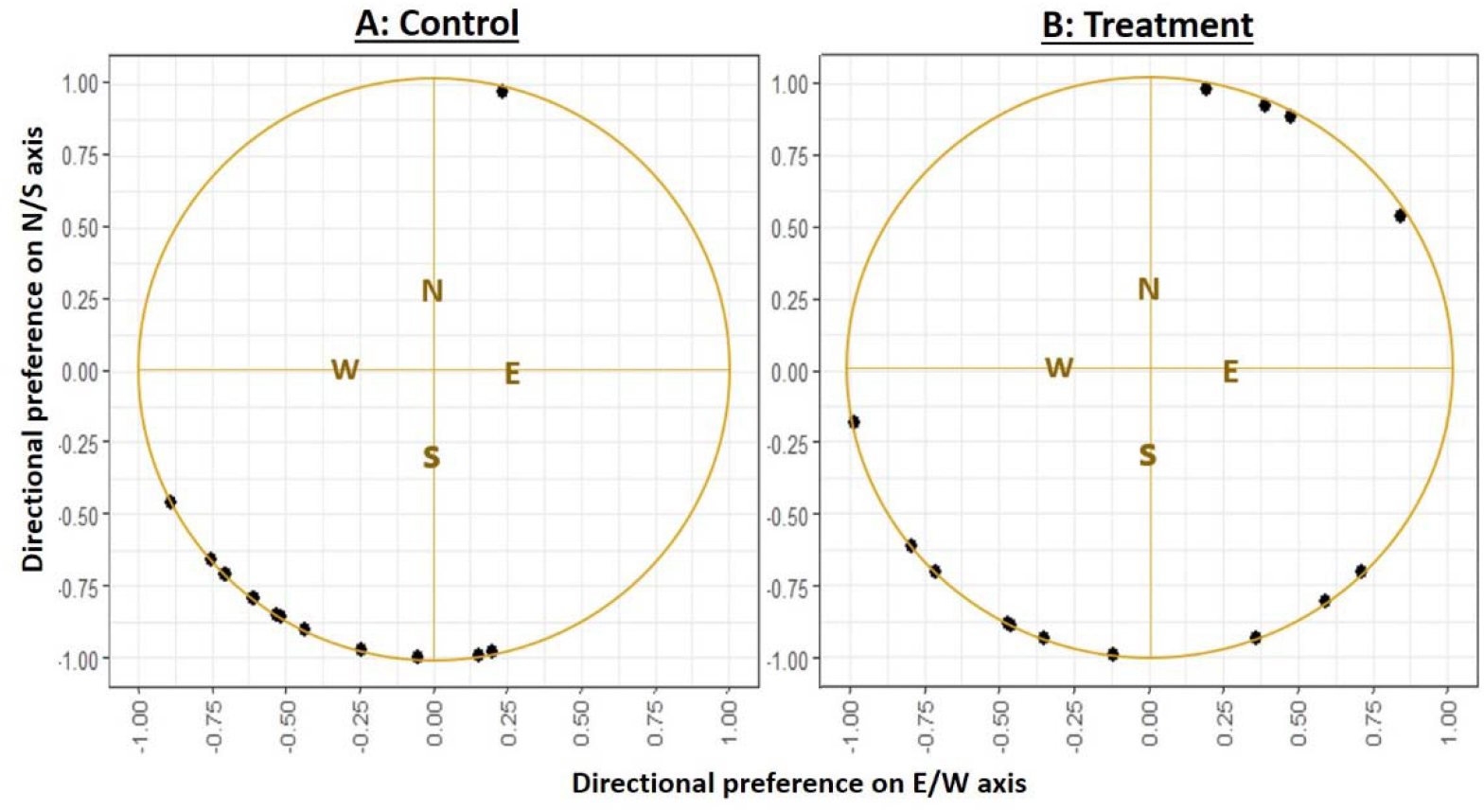
Orientation of treated and untreated reed warblers: Each dot represents the average hop azimuth of an individual in an Emlen funnel. Scales are projections of azimuths on x/y axes in the trigonometric unit circle (see Materials and Methods). On the north/south axis, positive values indicate a northward tendency and negative values indicate a southward tendency. On the east/west, positive values indicate east, while negative values indicate west.

**Figure 2:**
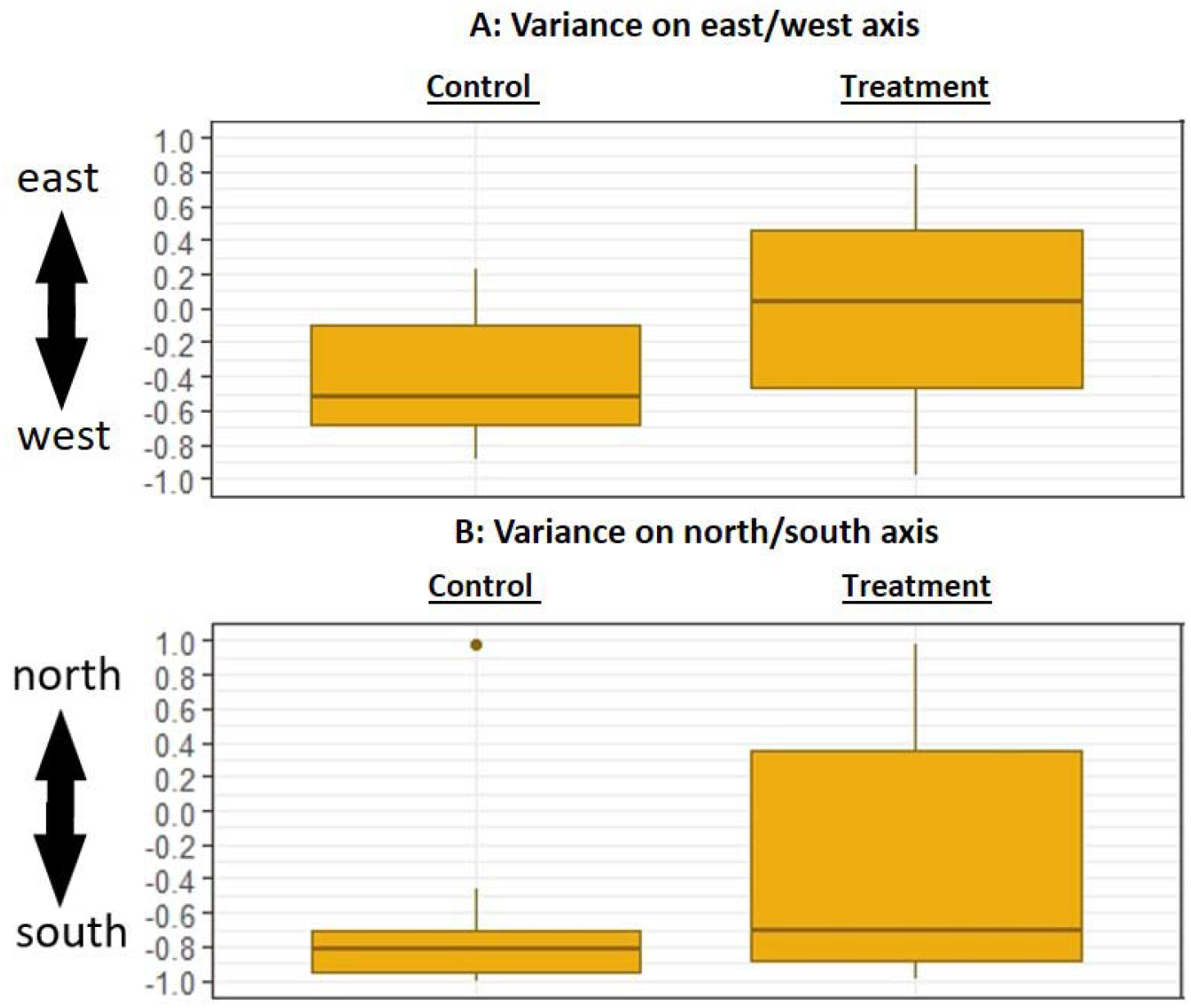
Variance of directional preference: Differences in the variance of directional tendency between treatment and control groups. Scales are projections of azimuths on x/y axes in the trigonometric unit circle (see Materials and Methods).

The results obtained from implementation of the well-established Emlen protocol indicate that Enrofloxacin affects orientation-related behavior during migration restlessness in a night-migrating passerine. This is the first documentation of such an effect. As seen in Figure 1, treatment group graphs are similar to control graphs, with the exception that in the first, individual azimuths are more dispersed over the diagram. Significant orientation is a function of low directional variance. The significant difference in directional variance between the experimental groups is exactly what would be expected if antibiotics are detrimental for orientation. This difference is significant regardless of the lack of significant orientation of the treatment group. The size and shape of the funnel do not allow birds to do much more than take a small skip towards the direction in which they have decided to go, meaning that each hop should represent the directional decision made prior to jumping. This narrows the window of antibiotic effects further, to the decision-making process itself. Directional decisions during migration depend on multiple internal and external factors [17]. Emlen funnels allow us to examine this process in a highly controlled setting. According to widely used, standard techniques, in our setup tested birds could only rely on magnetic information for the process of directional decision making, as they had no access to other celestial cues. This narrows our window of effect further still. The choice of specimens was such that only migrants at late stages of preparation to migrate were tested. From these, only adjusted, heavily fueled individuals were tested. A three-day stay in a cage, including treatment and handling prior to migration could have affected the resulting patterns in various ways. But in this respect, individuals from both groups underwent exactly the same procedure. This leaves us with the potential physical effects of the antibiotic, Enrofloxacin, as the internal cause of the differences between groups. The amount needed to cause any effect to vertebrate cells is two orders of magnitude larger than the bactericidal, therapeutic amount [18]. This means that any disruption of the directional, decision-making process caused by our treatment should have been mediated by a bacterial factor, assuming, as discussed above, that the only external directional stimuli to which the birds were exposed was magnetic. Namely, the process of directional decision making involves (at least in part) bacterial factors. We regard these results as experimental support for the symbiotic magnetic sensing hypothesis.

Today, many processes in multicellular organisms are found to include bacterial involvement [19–21]. Thus, the conclusions from this experiment are not surprising. An immediate conclusion would therefore be that antibiotic pollution should be considered a global concern not only in the context of pathogen resistance and human health [22] but also for processes involving magnetic field sensing such as migration in the air, on land, or at sea, pollination, habitat choice, etc.

We expected the directional tendency in spring to be northwards, according to the migration route of the species in the region [13]. The resulting south-western tendency could be explained in several ways. Prior work with the study species has shown it to demonstrate axial behavior, meaning that part of the population (up to 55%) shows reverse directionality with regard to migration route [23]. Furthermore, “reverse migration” is a known phenomenon, which has been related to stress, late night migration [24] and the lack of exposure to celestial information and other directionally significant environmental factors for a long duration prior to, and during the test [17,25,26]. The issue of exposure to celestial cues during Emlen tests is debatable, and there are groups working on both methods [17,5]. We aimed to isolate the magnetic factor of orientation, so obstruction of celestial cues was important. Regardless of the reason for the seasonally inappropriate directionality, we show significant directional tendencies in control groups which were absent in treatment groups. Most importantly, we show a significant increase in directional variance in the antibiotic treatment group. Considering all the above, this trend could indicate bacterial involvement in navigation-related processes in a passerine, specifically magnetic sensing.

## Acknowledgments

We would like to thank Shay Agmon of the Hula ringing station and the Hula research center staff for their assistance throughout the project.

